# Multiple axes of visual system diversity in Ithomiini, an ecologically diverse tribe of mimetic butterflies

**DOI:** 10.1101/2023.07.18.549450

**Authors:** J. Benito Wainwright, Corin Schofield, Max Conway, Daniel Phillips, Elizabeth Martin-Silverstone, Emelie A. Brodrick, Francesco Cicconardi, Martin J. How, Nicholas W. Roberts, Stephen H. Montgomery

**Affiliations:** School of Biological Sciences, University of Bristol, 24 Tyndall Avenue, Bristol, BS8 1TQ, UK; Bristol Palaeobiology Group, School of Earth Sciences, University of Bristol, 24 Tyndall Avenue, Bristol, BS8 1TQ; Living Systems Institute, University of Exeter, Stocker Road, Exeter, EX4 4QD

**Keywords:** Allometric scaling, apposition compound eye, Lepidoptera, neuroecology, optic lobe, visual ecology

## Abstract

The striking structural variation seen in arthropod visual systems can be explained by the overall quantity and spatio-temporal structure of light within habitats coupled with developmental and physiological constraints. However, little is currently known about how fine-scale variation in visual structures arise across shorter evolutionary and ecological scales. In this study, we characterise patterns of interspecific (between species), intraspecific (between sexes) and intraindividual (between eye regions) variation in the visual system of four ithomiine butterfly species. These species are part of a diverse 26-Myr-old Neotropical radiation where changes in mimetic colouration are associated with fine-scale shifts in ecology, such as microhabitat preference. By using a combination of selection analyses on visual opsin sequences, in-vivo ophthalmoscopy, micro-computed tomography (micro-CT), immunohistochemistry, confocal microscopy, and neural tracing, we quantify and describe physiological, anatomical, and molecular traits involved in visual processing. Using these data, we provide evidence of substantial variation within the visual systems of Ithomiini, including: i) relaxed selection on visual opsins, perhaps mediated by habitat preference, ii) interspecific shifts in visual system physiology and anatomy, and iii) extensive sexual dimorphism, including the complete absence of a butterfly-specific optic neuropil in the males of some species. We conclude that considerable visual system variation can exist within diverse insect radiations, hinting at the evolutionary lability of these systems to rapidly develop specialisations to distinct visual ecologies, with selection acting at both the perceptual, processing, and molecular level.

**Summary statement:** Physiological, anatomical, and molecular evidence of extensive visual system variation within a diverse butterfly radiation, hinting at the lability of visual systems to evolve specialisations to distinct visual environments.

## Introduction

The diversity of animal visual specialisations reflects their central role in key behaviours such as foraging, navigation, communication, and predator avoidance (Cronin *et al*. 2014; Endler *et al*. 2005). As the maintenance of complex visual adaptations incurs significant energetic costs (Land & Nilsson, 2012; Laughlin *et al*. 1998; Moran *et al*. 2015; Niven *et al*. 2007; Niven & Laughlin, 2008) divergence in specialization at any functional, physiological, anatomical or molecular level is likely the result of ecological selection pressures (e.g. Gonzalez-Bellido *et al*. 2011; Hofmann *et al*. 2009; Huber *et al*. 1997; Scales & Butler, 2016; Sugawara *et al*. 2005; Zhao *et al*. 2009). However, such specializations are also limited by a number of trade-offs including evolutionary history, body size and other allometric and physiological constraints (Land, 1997; Land & Nilsson, 2012; Warrant & McIntyre, 1993).

Understanding how an animal’s visual ecology interacts with these constraints to shape variation in visual systems is a central question to sensory and neuroecology (Cronin *et al*. 2014; Lythgoe, 1979; Striedter, 2005).

Visual system diversity is greatest within the Arthropoda (Osorio *et al*. 1995; Stansbury & Moczek, 2013; Yilmaz *et al*. 2022). This diversity is perhaps best reflected by their colour vision which is acquired via the tuning of visual pigments found within photoreceptor cells. These pigments consist of a retinal-based chromophore attached to an opsin protein, where the sequence of critical amino acid residues within the chromophore binding pocket of the opsin can shift the wavelength sensitivity which varies between species with different visual ecologies (Fain *et al*. 2010; Feuda *et al*. 2016; Frentiu *et al*. 2007; Terakita, 2005). In apposition compound eyes, each facet typically projects light along a single rhabdom within an ommatidium, separated from adjacent ommatidia by light-absorbing pigments (Land, 1989). However, anatomical, and physiological parameters such as total eye size, shape, lens diameter, acuity, sensitivity, number of ommatidia, and pupillary response vary, even between closely related species (Bartholomée *et al*. 2023; Greiner, 2006; Greiner *et al*. 2004a; Land, 1989; Narendra *et al*. 2013; Scales & Butler, 2016; Somanathan *et al*. 2009; Warrant, 2001). For example, shifts to a nocturnal lifestyle have led to larger ommatidial facets and rhabdom diameters in the apposition eye morphology of the sweat bee, *Megalopta genalis,* compared to its close relatives (Greiner *et al*. 2004a; Warrant, 2017). Microhabitat partitioning within communities of Hawaiian damselflies and hemiboreal bumblebees has also resulted in rapid shifts in eye sensitivity at a much finer ecological scale (Bartholomée *et al*. 2023; Scales & Butler, 2016). Spatial variation in light abundance and spectral composition within habitats has also promoted compound eye regionalisation whereby different eye regions are fine-tuned to optimally receive ecologically relevant stimuli (e.g. Arikawa *et al*. 2009; Lehrer, 1998; Labhart & Meyer, 1999; Meyer & Labhart, 1992; Nilsson & Smolka, 2021; Nilsson *et al*. 2022; Stavenga, 1992; Stavenga *et al*. 2001; White *et al*. 2003; Zufall *et al*. 1989). For example, the dorsal rim area (DRA) of many insects is specialised for polarization vision, useful for orientation and navigation (Dacke *et al*. 2002; Mappes & Homberg, 2004; Reppert *et al*. 2004; Sauman *et al*. 2005; Stalleicken *et al*. 2006). The fine structure and function of the primary insect visual neuropils can also differ (Sinakevitch *et al*. 2003). While the general structure of optic lobes is largely conserved, typically subdivided into four main synapse-dense regions (the lamina, medulla, lobula and the lobula plate) (Fig. 1a; Strausfeld & Nässel, 1980), the way they have evolved to process different forms of visual information again reflects their visual environment and evolutionary history. For example, neural adaptations for spatial and temporal summation within the lamina are repeatedly associated with shifts towards nocturnal diel patterns (Greiner *et al*. 2004b; Warrant *et al*. 2004, Stöckl *et al*. 2016b; 2020).

**Fig. 1:**
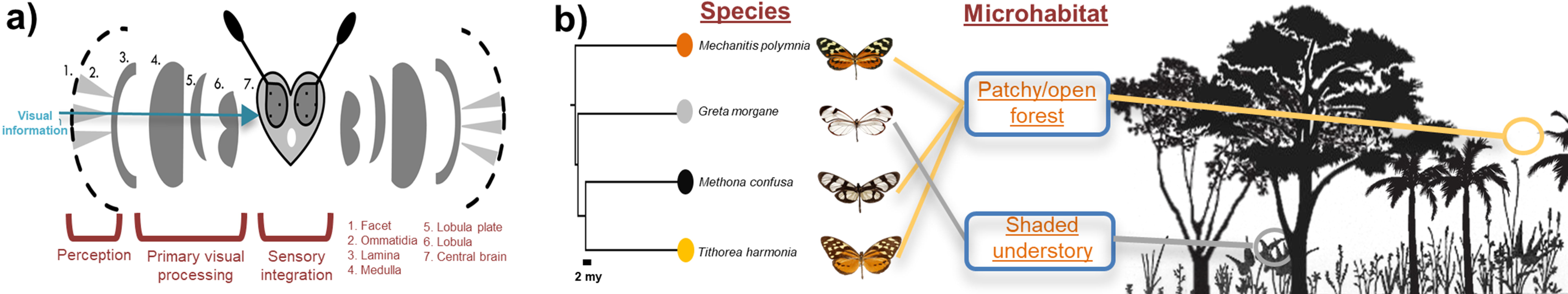
A**n overview of butterfly visual systems and ecology of the four ithomiine study species.** (a) Schematic diagram of a butterfly head representing the different levels of sensory organisation within the visual system. (b) Microhabitat segregation in ithomiines coupled with a pruned molecular phylogeny calibrated by Chazot *et al*. (2019) to show the evolutionary relationships between the four study species.

Understanding how visual systems evolve in response to novel habitats requires comparative study systems of species with diverse and well documented ecologies. Lepidopterans are one such group, occupying a wide range of light environments, relying heavily on the use of visual cues and signals for a variety of behavioural tasks, and exhibiting considerable levels of investment in specialized visual system physiology and anatomy (Bergman *et al*. 2021; Briscoe & Chittka, 2001; Couto *et al*. 2020; Nilsson *et al*. 1988; Stavenga & Arikawa, 2006; St[ckl *et al*. 2016a,b). Like other arthropods, lepidopteran species that occupy high-light intensity environments tend to have larger optic lobe neuropils than species found in low-light habitats, or nocturnal conditions (Couto *et al*. 2020; Heinze & Reppert, 2012; Montgomery & Ott, 2015; Montgomery & Merrill, 2017; Wainwright & Montgomery, 2022). The opposite appears true for relative eye size, where nocturnal species have evolved larger facets and wider and longer rhabdoms to maximise light collection and absorption (Frederiksen & Warrant, 2008; Yack *et al*. 2007). However, in butterflies there is also some evidence that visual systems vary across more subtle ecological differences, and readily adapt over short evolutionary scales. For example, a comparison of two *Boloria* butterfly species identified larger eyes and facets in the frontal eye region in *B. aquilonaris* whose habitat is naturally fragmented relative to *B. eunomia*, suggesting that differences in eye morphology can evolve remarkably rapidly to ecological challenges (Turlure *et al*. 2016). Consistent patterns of neural divergence in response to ecological preference shifts have also been observed between parapatric *Heliconius* butterfly species, which also display eye structural variation, separated across continuous environmental gradients within tropical forests (Hebberecht *et al*. 2023; Montgomery & Merrill, 2017; Montgomery *et al*. 2021; Seymoure *et al*. 2015).

To better explore how visual systems evolve in response to subtle environmental changes we present here a detailed study of interspecific and intraspecific differences in both eye and brain structure, across representatives of a diverse 26-Myr-old tribe of Neotropical butterflies, the Ithomiini (Nymphalidae: Danainae) (Chazot *et al*. 2019). Historically, ithomiines have been primarily studied for their Müllerian mimicry rings (also referred to as mimicry complexes) where multiple species have evolved convergent wing colours, patterns, and morphologies to amplify their aposematic signal to predators within sympatric communities (Bates, 1862; Beccaloni, 1997; Elias & Joron, 2015; Hill, 2021; Müller, 1879). Critically, for mimicry to be effective, co-mimics must signal to the same predators, whose distribution and abundance varies across forest habitats (Gompert *et al*. 2011; Willmott *et al*. 2017). As a result, data on forest structure suggest that mimicry rings are segregated across microhabitats, which likely expose closely related species from different mimicry rings to divergent visual environments (Beccaloni, 1997; Elias *et al*. 2008; Hill, 2010; Wainwright & Montgomery, 2022; Willmott *et al*. 2017). Typically, colourful ithomiine mimics are found in more open forest compared with cryptic, transparent-winged ithomiines which reside in shaded inner- forest (Elias *et al*. 2008; Pliske, 1975), and these habitat shifts have been linked to variation in investment in sensory neuropils (Montgomery & Ott 2015; Wainwright & Montgomery, 2022). In addition, while ithomiines are generally not sexually dichromatic, evidence of dimorphism in olfactory and visual processing centres might hint at the possibility of additional sex-specific patterns of visual investment (Montgomery & Ott 2015; Morris *et al*. 2021; Pliske, 1975; Wainwright & Montgomery, 2022).

Here, we focus on representatives of four subtribes within the ithomiine radiation: *Greta morgane* (subtribe: Godyridina), *Mechanitis polymnia* (subtribe: Mechanitina), *Methona confusa* (subtribe: Methonina) and *Tithorea harmonia* (subtribe: Tithoreina), hereafter referred to by their genus name for brevity (Fig. 1b; Chazot *et al*. 2019; de Silva *et al*. 2016). *Mechanitis, Methona* and *Tithorea* have converged on an open forest microhabitat, flying at high elevations (Elias *et al*. 2008), whereas the transparent *Greta* flies lower down in shaded forest understory (Fig. 1b; Elias & Joron, 2015; Willmott & Mallet, 2004). By combining results from selection analyses on opsin sequences from recent genomic data, ophthalmoscopy, micro-CT, immunohistochemistry, and neural tracing, we describe and quantify variation for several aspects of the ithomiine visual system. Our data reveal multiple examples of anatomical, physiological, and molecular variation within and between these four ithomiine species, highlighting potential adaptations for optimising visual perception and processing during microhabitat niche partitioning.

## Methods

### 1. Molecular analyses of visual opsins

#### i. Opsin gene alignment and data mining

Opsin sequences for *Danaus plexippus* (Nymphalidae: Danainae), the most closely related species to the Ithomiini for which all three visual opsins (ultraviolet, blue and long-wavelength) are publicly available on GenBank, were used as query sequences. These sequences were utilised for finding homologous opsins within recently assembled genomes of each of four ithomiine species (Cicconardi *et al*. in prep) using Exonerate v2.2.0 with ‘-model protein2genome -percent 50’ options (Slater & Birney, 2005; Zhan & Reppert, 2013). To increase the power of our comparative analyses, we used GenBank deposited ultraviolet (UV), blue (B) and long-wavelength (LW) opsin sequences from 13 additional nymphalid butterfly species with well-established ecologies (all sequences used in our analyses, including accession IDs, can be found in Table S1).

To confirm functional opsin number for each ithomiine species, the transmembrane structure of each opsin sequence was predicted using Phobius, implemented through Protter (Kall *et al*. 2007; Omasits *et al*. 2014), a webserver-based tool for making protein structural predictions. Any opsin sequences predicted to have six or seven transmembrane domains were considered putative functional opsins. Rh7 opsins were also surveyed, however this gene family was not included in downstream analyses due to a poor abundance of other available nymphalid Rh7 sequences, as previously noted by Sondhi *et al*. (2021). The 3D protein structure for the *Greta* and *Tithorea* UV opsins were also modelled using a homology-based approach in Swiss-Model (Waterhouse *et al*. 2018). The jumping spider Rhodopsin-1 (9i9k.1.A) was used as a template because this had the highest identity score and coverage (*Greta*: GMQE = 0.67, identity = 37.39; *Tithorea*: GMQE = 0.68, identity = 38.34).

#### ii. Gene tree estimation

Gene sequences for each opsin family (UV, B, LW) were aligned separately using MACSE v2 (Ranwez *et al*. 2018), a tool which accounts for underlying codon structure when aligning protein-coding nucleotide sequences. Aligned sequences were manually cleaned and trimmed in SeaView v5 (Gouy *et al*. 2021) to ensure the alignment contained no stop or incomplete codons. The final alignments all contained approximately 1100 nucleotide bases with 370 amino acids which falls within the range of previously characterised lepidopteran opsin sequences (White *et al*. 2003; Zhan & Reppert, 2012). IQ-TREE (multi-core v1.6.12) was then used to build a maximum likelihood (ML) nucleotide gene tree for each opsin from these alignments (iqtree -s alignment_name.fasta -st DNA -bb 10000 -nt AUTO -alrt 1000) using ModelFinder and ultrafast bootstrap (Fig. 2a; Hoang *et al*. 2018; Kalyaanamoorthy *et al*. 2017; Trifinopoulos *et al*. 2016).

**Fig. 2:**
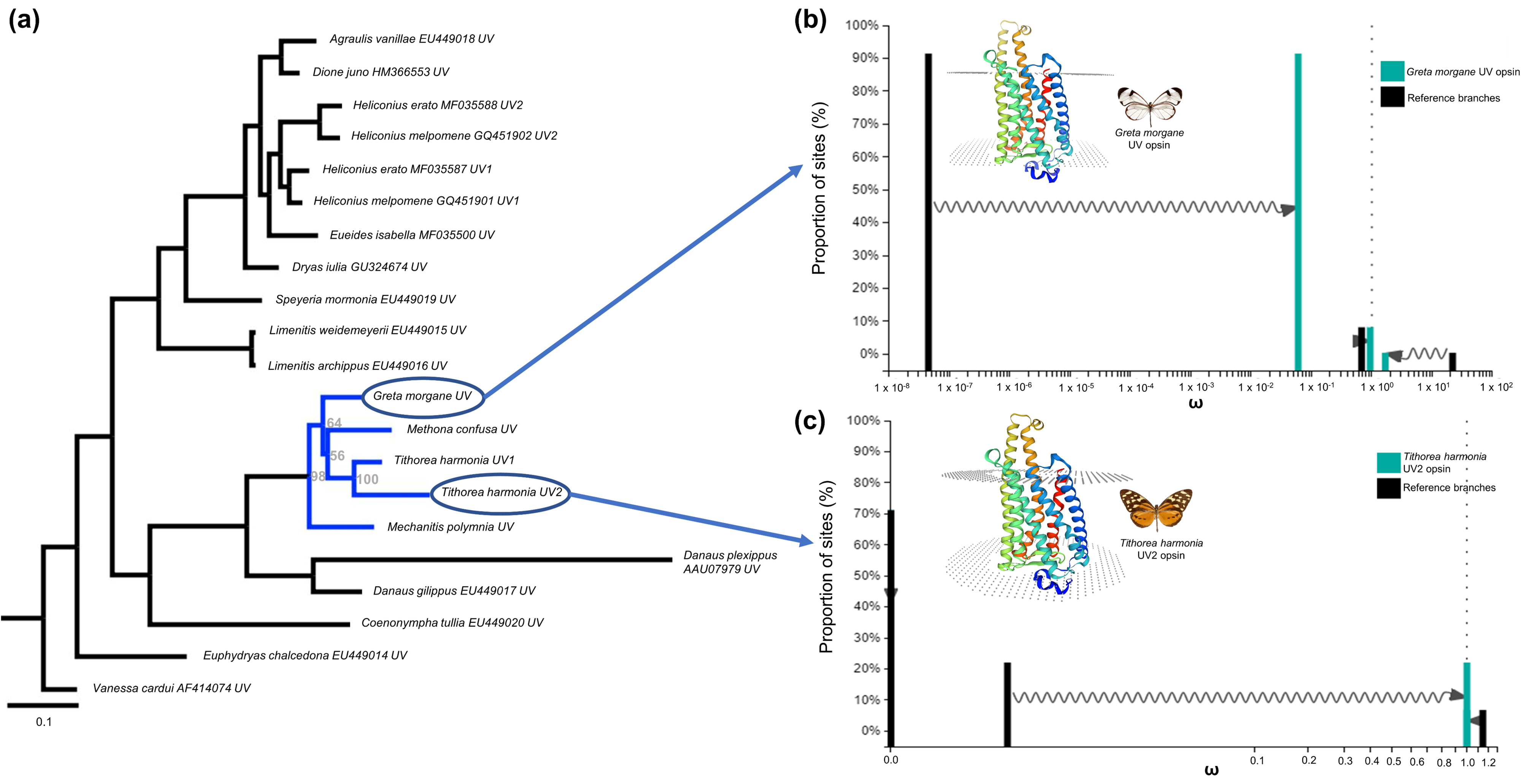
S**i**gnatures **of relaxed selection in ithomiine UV visual opsins.** (a) Maximum likelihood nucleotide gene tree created using IQ-TREE which includes all UV opsin sequences used in our analyses. Ithomiine branches are highlighted in blue for which the ultrafast bootstrap values calculated from 10000 replicates are displayed at the nodes. (b) Comparison of ω distributions between the *Greta morgane* UV test branch (turquoise) and all other reference branches under the alternative model of relaxed selection. The branch tips leading to the duplicated UV2 opsins of *Heliconius melpomene*and *Tithorea harmonia* were left as unspecified in the RELAX analysis. (c) Comparison of ω distributions from the newly discovered UV2 opsin of *Tithorea harmonia* under the alternative relaxed selection model when compared to all other reference branches. Branches leading to the *Greta morgana* UV and *Heliconius melpomene* UV2 opsins were left unspecified. In both (b) and (c), sites under purifying (ω > 1) or positive (ω < 1) selection in the reference move towards neutrality in the test branch (ω = 1). Arrows indicate that when comparing sites in the prespecified test and reference branches, all three ω distributions used in the BS-REL model (corresponding to sites under strong purifying, weakly purifying, and positive selection) shift closer towards neutrality in the test branches. Superimposed on both (b) and (c) is a 3D protein model of the *Greta morgane* UV and *Tithorea harmonia* UV opsin respectively, created using Swiss-Model (Waterhouse *et al*. 2018) with the jumping spider rhodopsin-1 as the template. Prediction of alpha helices are shown in different colours and areas between the top and bottom dotted layers indicate transmembrane predictions.

#### iii. Selection analyses

We tested whether an ecological shift from open to shaded forest was associated with relaxed or intensifying selection on the visual opsins of *Greta* by applying RELAX implemented in HyPhy, accessed via the Datamonkey webserver (Kosakovsky Pond *et al*. 2020; Muse *et al*. 2018; Wertheim *et al*. 2015). When given prespecified test (*T*) and reference (*R*) branches from the gene tree, RELAX estimates the rate of nonsynonymous to synonymous nucleotide base substitutions (d_N_/d_S_, or ω) among three rate categories using a BS-REL (branch site random effects likelihood) model (Kosakovsky Pond *et al*. 2011) before raising these values to the parameter k (ω^k^) in *T*, which estimates the strength of selection at *T* relative to *R*. When k = 1, as is the case in the null model, the ω distributions of *T* and *R* are the same. If k < 1, *T* is under relaxed selection relative to *R*, whereas if k > 1, *T* is under intensified selection relative to *R* (Wertheim *et al*. 2015). Null and alternative models are then compared with likelihood ratio tests using a χ^2^ distribution. In our analyses, the *Greta* branch tip was specified as *T* for each opsin class with all other branches being assigned to *R,* with the exception of the UV opsin analysis where duplicated opsins were included, with these portions of the tree being left as unspecified branches. Additional tests were performed to compare patterns of selection following gene duplications, where the branch stemming from the duplicated node were specified as *T* and all other branches assigned as *R*.

### 2. Animals for physiology and anatomy

Butterflies used in physiological and anatomical comparisons were purchased as pupae or as live adults from The Entomologist Ltd (*Mechanitis, Methona* and *Tithorea*) and Tropical Butterflies UK (Edu-Sci Ltd) (*Greta*) respectively. Individuals were reared in 2.2 x 1.8 x 1.8m cages kept at 28-35°C and 80% relative humidity at the University of Bristol’s Old Park Hill Greenhouse facility and were regularly fed 30% sucrose solution. Adults were marked and sexed immediately post-eclosion and matured for a minimum of three days before being sampled.

### 3. Eye physiology comparisons

#### i. Ophthalmoscopy

Interspecific and intraspecific variation in eye physiology were explored using a custom-built ophthalmoscope (Fig. S1), based on the system described by Brodrick *et al*. (2020). It consisted of a UI-3590CP-C-HQ-R2 camera with a CMOS colour sensor (Imaging Development Systems, Germany), 10x 0.25 NA objective lens (Plan N, Olympus, Tokyo, Japan), beam-splitter (Thorlabs, Newton, MA) and a portable fibre-coupled broadband (470- 850 nm) light emitting diode (LED, MBB1F1 Thorlabs, Munich, Germany) which was projected along an optical fibre, providing co-axial illumination of the eye. The camera was connected to a computer with the uEye Cockpit program (part of the IDS Software Suite 4.95) installed for capturing and recording images and videos.

Twelve live individuals of each of the four species were used in the investigation, with approximately equal numbers of males and females (Table S2). Intact butterflies were mounted in slotted plastic tubes and further immobilised using plasticine before being suspended at the centre of rotation of a Newport M-MGM80MS 90° rotation goniometric cradle by attaching the plastic tube to the arm of a micromanipulator (Fig. S1). Butterflies were then oriented to set the eye’s direction of view. The ophthalmoscope was adjusted to focus on a region of the cornea where the optical axes of several ommatidia are directly facing the objective lens, therefore viewing the individual’s pseudopupil. After dark adapting for as briefly as 10 seconds, the LED can be switched on to reveal the eyeshine (or luminous pseudopupil) created by unabsorbed light reflecting off a mirror-like tracheolar tapetum at the proximal end of each rhabdom (Franceschini & Kirschfeld, 1971; Stavenga, 1979, 2002). Unabsorbed reflected light leaves the eyes, referred to as eyeshine, the colour and heterogeneity of which varies hugely between butterfly species (Belušič *et al*. 2021; Briscoe & Bernard, 2005; Stavenga *et al*. 2001). As expected, the intensity of this eyeshine diminished within seconds of illumination due to the intracellular migration of pigment granules which move from the soma towards the rhabdomeres, preventing light from reaching the tapetum (Qiu *et al*. 2002; Stavenga *et al*. 1977). This is known hereafter as the pupillary response. We used the duration of the pupillary response to assay how rapidly eyes physiologically respond to sudden changes in their light environment.

Video recordings of the eyeshine were taken in the dorsal, frontal, and ventral eye regions (achieved by rotation of the goniometric cradle) after ten minutes of dark adaptation under standard laboratory conditions (21°C). When video recordings were made in the frontal eye region, individuals were also dark adapted for 0.5, 1, 5 and 20 minutes, in a randomised order, to confirm whether any interspecific patterns are still observed for different lengths of dark adaptation. We also recorded whether the rate of habituation to a flashing light stimulus differed between species by switching the LED light source on and off at ten second intervals for a period of five minutes. Here, our intention was to simulate changes in light conditions as individuals transition between microhabitats. Videos were imported into FIJI/ImageJ (Schindelin *et al*. 2012) where response time was visualised and estimated using the View5D plugin (Fig. 3a). Reflecting facets were counted and categorized as being red or yellow, where red reflecting ommatidia indicate the presence of long-wavelength shifted photoreceptors. Using this, the ratio of yellow:red ommatidia was also calculated for the frontal region of each individual.

**Fig. 3:**
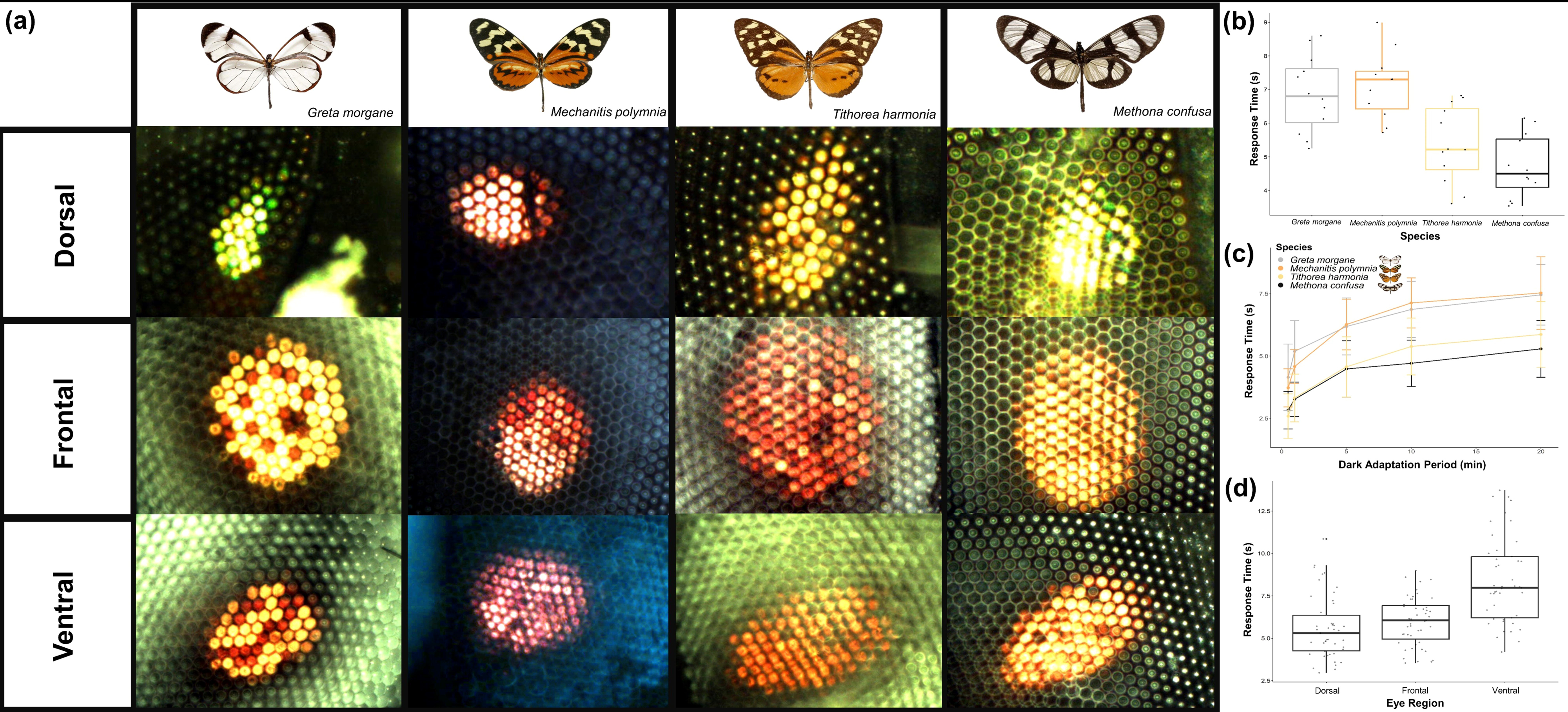
I**t**homiine **eyeshine variation** (n = 48, 54.17% female). (a) Ophthalmoscope images of the dorsal, frontal, and ventral luminous pseudopupil within a single individual of all four study species after ten minutes of dark adaptation. (b) Interspecific differences in frontal pupillary response time. (c) Frontal pupillary response time after different lengths of dark adaptation. Points show means for each species (n = 12 per species, 50-58% female) with error bars representing standard deviation. (d) Intraindividual differences in pupillary response time for the dorsal, frontal, and ventral eye regions. For all boxplots, medians (thick horizontal bars), interquartile ranges (boxes), values within 1.5 interquartile ranges of the box edges (whiskers) and possible outliers (data points outside the whiskers) are plotted.

Once eye physiological data had been collected, half of the individuals from the eyeshine study were sampled for micro-computed tomography (micro-CT; see section 4). The second half of individuals from the eyeshine experiment were sampled for immunohistochemistry (see section 5).

#### ii. Statistical analysis

All statistical analyses were performed in R (R Core Team 2016). Following Shapiro-Wilk tests for normality and Bartlett’s test for homogeneity of variances, linear models were constructed, testing the effect of species, sex, and their interaction, on response time in the frontal region. Subsequent post-hoc comparisons were performed using the TukeyHSD function. Similar analyses were also performed to test for species and sex differences in response time in the habituation experiment. The ratios of yellow:red reflecting frontal ommatidia did not follow a normal distribution, so the non-parametric Kruskal-Wallis test was performed instead. Intraindividual differences in the number of reflecting ommatidia and response time between eye regions were analysed by building linear models with the mean fitted as a fixed effect, the null hypothesis being that the mean difference between eye regions is zero. For response time, species, sex, and their interaction were included as additional independent variables.

### 4. Micro-computed tomography

#### i Image acquisition

Heads were severed and placed directly into a chilled fixative (4% paraformaldehyde, 2% glutaraldehyde, 2% glucose) in 0.1 M sodium phosphate buffer solution (pH 7.4) for 24 hours under agitation. Heads were dehydrated in a graded ethanol series (10%, 30%, 50%, 70%, 90%, 95%, ten minutes each), before being stained with 1% iodine dissolved in 100% ethanol for three days under agitation to enhance X-ray absorption contrast (Smith *et al*. 2016; Swart *et al*. 2016). Samples were subsequently washed in 100% ethanol three times, each for 30 minutes and stored in ethanol at 4°C until ready for use. The body weight (g) of each individual was also recorded.

Tomographic images of the samples were taken at the University of Bristol’s X-ray tomography (XTM) facility using a Nikon XTH225ST scanner with a 180kV transmission target. Heads were typically imaged using x0.33 total magnification and a ∼2.2 µm effective pixel size (see Table S5 for full scanning parameters). 23 heads were scanned in total (n = 6 per species with the exception of *Tithorea harmonia* where one individual experienced bacterial growth and was thus removed from any subsequent analyses; see Table S3). Scan acquisition and reconstruction was achieved using Nikon CT Pro and the resulting TIF files were exported using VG Studio MAX (Fig. 4a). The resolution of the micro-CT scans allowed identification of the corneal lens of each ommatidium, as well as the area where the crystalline cones, and light sensitive rhabdoms are located (Fig. 4b-d).

**Fig. 4:**
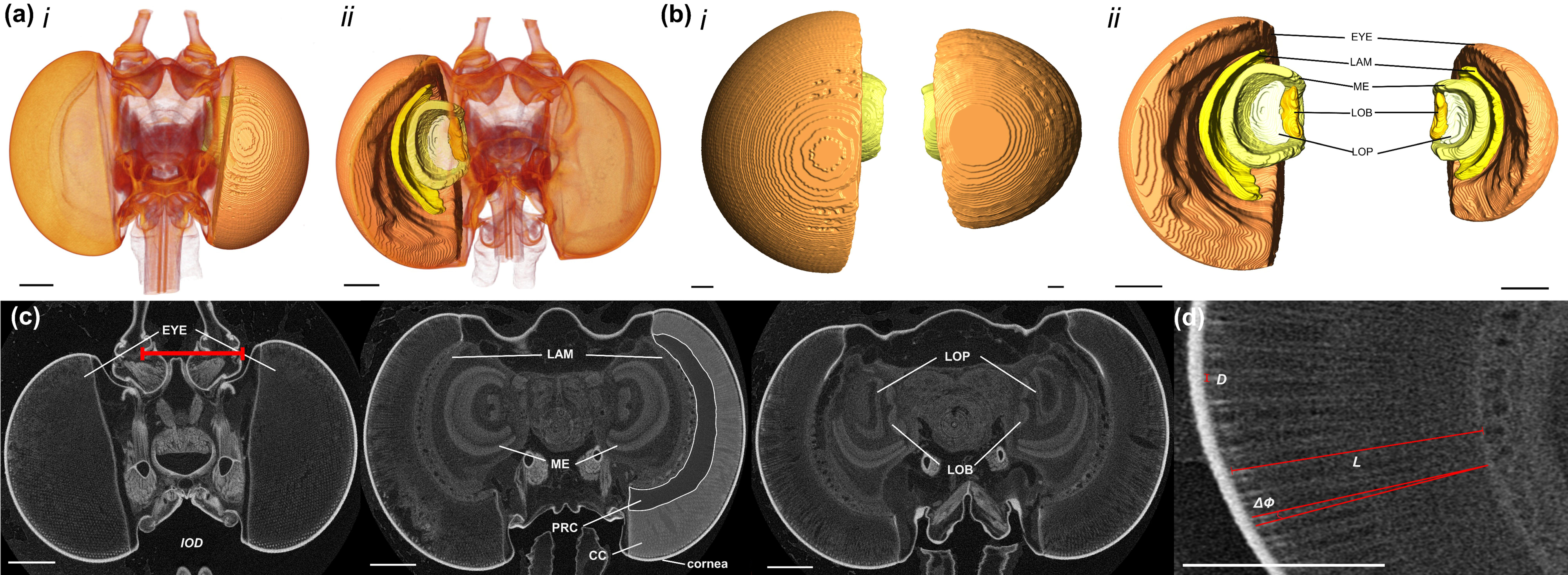
A**n eye and brain anatomical overview for the ithomiine species using micro-CT data** (n = 23, 52.17% female). (a) Volume rendered whole-head images from micro-CT scans of *Methona confusa* from the anterior (i) and posterior (ii) with surface reconstructions of the eye and the four primary optic neuropils of interest superimposed on the right. (b) Anterior (i) and posterior (ii) surface reconstructions of the eye (EYE), lamina (LAM), medulla (ME), lobula (LOB) and lobula plate (LOP) of *Methona confusa* (left) and *Greta morgane* (right), shown to scale. (c) Iodine-stained micro-CT X-ray tomography frontal sections of *Methona confusa* taken at progressively posterior positions through the head, moving from left to right. Additional labels are the cornea, the areas containing crystalline cones (CC) and photoreceptor cells (PRC) and an exemplification of our measure of interocular distance (*IOD*). (d) Further exemplifications of our measures of facet diameter (*D*), ommatidial length (*L*) and interommatidial angle (ΔФ) from sections of the same *Methona confusa* individual. Scale bars = 200 μm.

#### ii. Eye volumetric reconstructions

The resulting TIF stacks were compressed from 32 bit to 8 bit and cropped in FIJI/ImageJ (Schindelin *et al*. 2012). They were then loaded into Amira 3D 2021.2 (ThermoFisher Scientific, FEI Visualization Sciences Group). Label files were created for each individual using the *labelfield* module, and segmentation was performed by manually delineating the boundaries of the eye based on intensity contrasts (Fig. 4c,d). Features were labelled every 5- 10 slices before being interpolated so all intervening slices could be assigned to the structure of interest. Reconstructions were edited and smoothed before total volumes were extracted using the *measure statistics* module. The volume of each bilaterally paired structure was then multiplied by two and log_10_ transformed before any analyses.

#### iii. Eye anatomical measurements

Further anatomical measurements were made with the *3D Measurement* tool in Amira to manually estimate ommatidial length (*L,* µm), interommatidial angle (ΔΦ, °), and facet diameter (*D,* µm) (Fig. 4d,e). 5-10 measurements of *L*, ΔΦ and *D* were taken in each eye region with a mean calculated per region per individual. Eye regions were defined in the XY dimension with the upper, middle, and lower thirds of the eye comprising the dorsal, frontal, and ventral eye regions respectively. In practice, no measurements were taken in ambiguous regions and the distribution of dorsal and ventral measurements are skewed towards the extreme ends of those boundaries as a result. To minimise measurement error when estimating *D,* the width of five adjacent facets were measured before dividing by five. We also measured interocular distance (*IOD,* µm), defined as the minimum horizontal gap between the two compound eyes when viewed in the XY dimension, to provide an allometric control in our statistical analyses. As was the case for the volumetric reconstructions, all eye anatomical data were log_10_ transformed before any analyses.

#### iv. Statistical analysis

To test for allometric scaling differences, linear models were constructed where each measurement of interest was scaled against *IOD* with species, sex and their interaction included as additional independent variables. If significant species or sex effects were detected, post-hoc pairwise scaling comparisons were made by building standardised major axis regressions using the sma function in the *smatr* package (Warton *et al*. 2012). This function tests for group level deviation from a common scaling relationship between two traits, modelled as log y = βlog x + α. We first tested for conservation in the allometric slope (β), deviation from which would suggest an effect of species or sex in how each dependent variable interacts with *IOD*. If equal slopes were found, the presence of a ‘grade shift’ (change in α) along the y-axis was examined, a common indicator of adaptive sensory divergence (Farnworth & Montgomery, 2022; Kruska, 2005; Montgomery *et al*. 2016; Sylvester *et al*. 2011). Facet diameter (*D*) did not scale with *IOD* (or central brain or body size) suggesting that any differences between species occur independently of allometric scaling. We therefore tested for differences using the absolute, untransformed *D* values.

Intraindividual differences in eye structure between dorsal, frontal, and ventral eye regions were analysed by applying the same statistical methods used for the eyeshine data, described above.

### 5. Neuroanatomical comparisons

#### i. Immunohistochemistry

Brain dissections were conducted under HEPES-buffered saline (HBS; 150mM NaCl; 5mM CaCl_2_, 25mM sucrose; 10mM HEPES; pH 7.4) with brains subsequently fixed in zinc formaldehyde solution (ZnFA; 0.25% [18.4mM] ZnCl_2_; 0.788% [135mM] NaCl; 1.2% [35mM] sucrose, 1% formaldehyde) for 16-20 hours under agitation, as in Ott (2008). Brains were washed three times in HBS and placed in Dent’s solution (80% methanol/20% DMSO) for ∼two hours under agitation. They were then placed in 100% methanol for one hour at room temperature before being transferred to fresh methanol and stored at -20 °C until later use.

Rehydrated brains were embedded in agarose gel (5% UltraPure agarose power in deionised water) and sectioned using a vibrating blade microtome with an 80 µm interval (Leica VT1200, Wetzlar, Germany). Sections were then washed six times in PBS-TX (0.5% Triton X-100 detergent in phosphate-buffered saline), five minutes each, before being incubated in 5% normal goat serum dissolved in PBS-TX (NGS-PBS-TX; NGS; New England BioLabs, Hitchin, Hertfordshire, UK) for a minimum of one hour at room temperature. Samples were then stained with anti-allatostatin (Antibody 5F10; Developmental Studies Hybridoma Bank, University of Iowa, Iowa City, IA, RRID: AB_528076) at a 1:100 dilution in fresh NGS-PBS-TX and incubated for 24 hours at room temperature under agitation. Allatostatin is a neuropeptide expressed in distinct cell types within the optic lobes of insects and crustaceans (Kreissl *et al*. 2010; Lin *et al*. 2021; Sivasubramanian & Sood, 2003) and provides additional anatomical detail of these brain regions. Slices then underwent 6 x 30 minute PBS-TX washes before introducing the secondary Cy2-conjugated anti-mouse antibody (Jackson ImmunoResearch Cat No. 115-225-146, RRID: AB_2307343, West Grove, PA) at a 1:100 dilution in fresh NGS-PBS-TX. Samples were left in the dark at room temperature for a further 24 hours, before undergoing 6 x 30 minute PBS-TX washes and added to 60% glycerol in PBS and left overnight. Slices were then transferred to 80% glycerol in PBS before mounting on glass slides with excess 80% glycerol solution under a cover slip, sealed with nail varnish.

#### ii. Dextran tracing and wholemount immunohistochemistry

In a separate sample of individuals, we performed *in vivo* dextran injections to explore neural connections between the optic lobe and structures within the central brain. Live butterflies were kept in custom-made slotted plastic holders and immobilised using dental wax. A small window was made in or just behind the compound eye, so part of the optic lobe was exposed. For some individuals, a window was made above the dorsal central brain instead to allow access to the mushroom bodies. A few crystals of dextran tetramethylrhodamine (Invitrogen^TM^ D1868,Fisher Scientific, Leicestershire, UK) were dissolved in bovine serum albumin (BSA, Merck Life Science UK Ltd, Gillingham, Dorset, UK) and placed on the tip of a horizontally pulled glass electrode (tip diameter ∼200 µm). Dextran was manually inserted into either both optic lobes or both mushroom bodies calyces under red-filtered light using a dissection microscope. Any remaining dextran on the brain surface was rinsed off with Ringer’s solution (Ringer; 150 mM NaCl, 3mM CaCl_2,_ 3mM KCl, 2mM MgCl_2,_ 10mM HEPES, 5mM glucose, 20mM NaOH, pH 6.9). The intact butterfly head was then left submerged in Ringer’s solution for a minimum of four hours at room temperature to allow the dextran to be transported to the terminal projection sites of affected neurons. The brain was then dissected and stored using the protocols described above.

To reveal neuropil structure, injected brains were stained as wholemounts using indirect immunolabelling against synapsin (Antibody 3C11; Developmental Studies Hybridoma Bank, University of Iowa, Iowa City, IA, RRID: AB_2315424), a conserved insect protein expressed at presynaptic regions (Brandt *et al*. 2005; Heinze & Reppert, 2011; Klagges *et al*. 1996; Montgomery & Ott, 2015). Brain samples were rehydrated in a decreasing methanol-Tris buffer dilution series (90%, 70%, 50%, 30%, 0%, pH 7.4) for ten minutes each. Samples were incubated in NGS (NGS; New England BioLabs, Hitchin, Hertfordshire, UK) diluted in 0.1 M PBS (pH 7.4) and 1% DMSO for two hours at room temperature, before the primary antibody was added at a 1:30 dilution with fresh NGS-PBSd and left at 4°C for 3.5 days under agitation. Non-bound primary antibody was removed after 3 x two hour PBSd washes before introducing the secondary Cy2-conjugated anti-mouse antibody at a 1:100 dilution in NGS-PBSd and left in the dark at 4°C for a further 2.5 days under agitation. Samples were then washed in glycerol diluted in 0.1 M Tris buffer (1% DMSO) in an increasing dilution series (1%, 2%, 4%, for two hours each, 8%, 15%, 30%, 50%, 60%, 70%, 80%, for one hour each) under agitation. Samples were dehydrated and washed in 100% ethanol three times, for 30 minutes each, before clarification by underlaying the ethanol with methyl salicylate.

#### iii. Confocal microscopy

All immunostained brains were imaged on a confocal laser-scanning microscope (Leica SP5- AOBS/SP5-II, Leica Microsystem, Mannheim, Germany) at the University of Bristol’s Wolfson Bioimaging Facility, using either a 10x 0.4 NA or 20x 0.7 NA objective lens (Leica Material Nos. 506285, 506513, Leica Microsystem). Single-capture images of wholemounts and sections were achieved with a 488 nm argon laser (20% intensity), and an x-y resolution of 512x512 pixels. For dextran-traced individuals, an additional 580 nm argon laser channel (20% intensity) was included and scanned sequentially with the 488 nm laser to visualise the passage of dextran dye against the synapsin-stained background.

#### iv Volumetric comparisons

Volumetric reconstructions of the four primary optic neuropils (lamina, medulla, lobula plate and lobula) were obtained from the micro-CT scans using the same methods described above (Fig. 7a). Paired neuropils were doubled and log_10_ transformed and analysed using the same statistical methods as the eye volumetric data. To test for associations between physically and functionally linked visual structures, a covariance matrix for the volumetric data was built by running multiple linear regressions with each structure of interest as a dependent variable and species as a random effect. We then tested for species and sex differences after accounting for these covariances by including species and sex as additional independent variables in these regressions.

## Results

### 1. Evidence of duplication and habitat-related relaxation of selection in UV opsins

We recovered single copies of UV, B and LW sensitive opsins in the genomes of all four ithomiine species, except for *Tithorea* where two putative full-length UV opsin sequences containing no stop codons were identified. Our selection analyses revealed the UV opsin of the shade dwelling *Greta* to show a significantly lower rate of nonsynonymous to synonymous nucleotide base substitutions relative to all assigned ‘open habitat’ reference branches (Fig. 2b; UV opsin, relaxation parameter, k = 0.17, χ2 test p = 0.016) when contrasted against the null model where all test and reference branches are assumed to be under the same selection pressures, indicating relaxed selection (see Methods). This null model was found to hold true when the same analyses were repeated on the B and LW opsin sequences (B opsin, k = 22.21, p = 0.314; LW opsin, k = 0.290, p = 0.174). The duplicated UV opsin of *Tithorea* also showed signatures of significant relaxed selection (Fig. 2c; k = 0.000, p < 0.001), potentially indicative of ongoing pseudogenization. Our results therefore may suggest shifts in selective regime operating at the molecular level, mediated by both habitat preference and gene duplication, which together might drive further variation in how visual systems respond to light.

### 2. The response to light varies across the eye and between species

We found significant variation in the retinal mosaic (Fig. 3) between species in the frontal region (yellow:Red reflecting frontal ommatidia, χ^2^ = 16.678, df =3, p = 0.001). *Greta, Mechanitis* and *Tithorea* all had distinguishable red and yellow reflecting facets, where *Tithorea* contained a greater proportion of red-reflecting ommatidia. *Methona* displayed more homogeneous orange eyeshine patterns (which were classified as red in the above analysis) (Fig. 3a, Table S4A). Therefore, all four species show evidence of a red-sensitive spectral channel. The number of shining ommatidia within each pseudopupil was greatest in the frontal region (dorso-frontal, t = 12.830, df = 37, p < 0.001; fronto-ventral, t = 3.540, df = 37, p = 0.003; Table S4D), suggesting that this is where spatial resolution is at its highest. The dorsal region contained the fewest shining facets.

When testing for differences in frontal pupillary response time (RT), we found evidence of significant interspecific variation in response time (F = 15.870, df = 3, p < 0.001) with no sex or interaction effects between species or sex (Fig. 3b, see Table S4B). Post-hoc analysis revealed that overall, *Greta* and *Mechanitis* had longer response times regardless of the length of dark adaptation (Fig. 3c, Table S4B). Response time decreased after five minutes of exposure to a flashing light stimulus (t = 2.354, df = 46, p = 0.011), but no differences in this habituation time were found between species, sex, or their interaction (see Table S4C).

Finally, the pupillary response of ventral ommatidia was significantly longer than those in the dorsal (t = 5.460, df = 46, p < 0.001) and frontal (t = 5.121, df = 46, p < 0.001) eye regions with no significant species, sex, or interaction effects (Fig. 3d). Significant species differences were only found when testing for dorso-frontal variation (F = 11.627, df = 3, p < 0.001) (Table S4E).

### 3. Eye structure varies between species, sexes, and eye regions

Micro-CT scans revealed quantifiable differences in overall eye size and structure between species (Fig. 4a,b). Each ommatidium within the apposition compound eye consists of a corneal lens and a deep crystalline cone which directs light through to the light-sensitive rhabdom where the photoreceptor cells are located (Fig. 1a, 4c,d). Relative eye volume varied significantly between species (F = 6.232, df = 3, p = 0.007) when scaled against *IOD*. Allometric analysis also revealed that these differences were a result of grade-shifts along the y-axis (Wald χ^2^ = 8.483, df = 3, p = 0.037, (Fig. 5a) with *Greta* investing more in eye volume than the similarly sized *Mechanitis* (Table S6), for example. Significant grade-shifts were also observed between *Methona* and *Tithorea*, with the latter sharing the same scaling relationship as its co-mimic, *Mechanitis* (Fig. 5a).

**Fig. 5:**
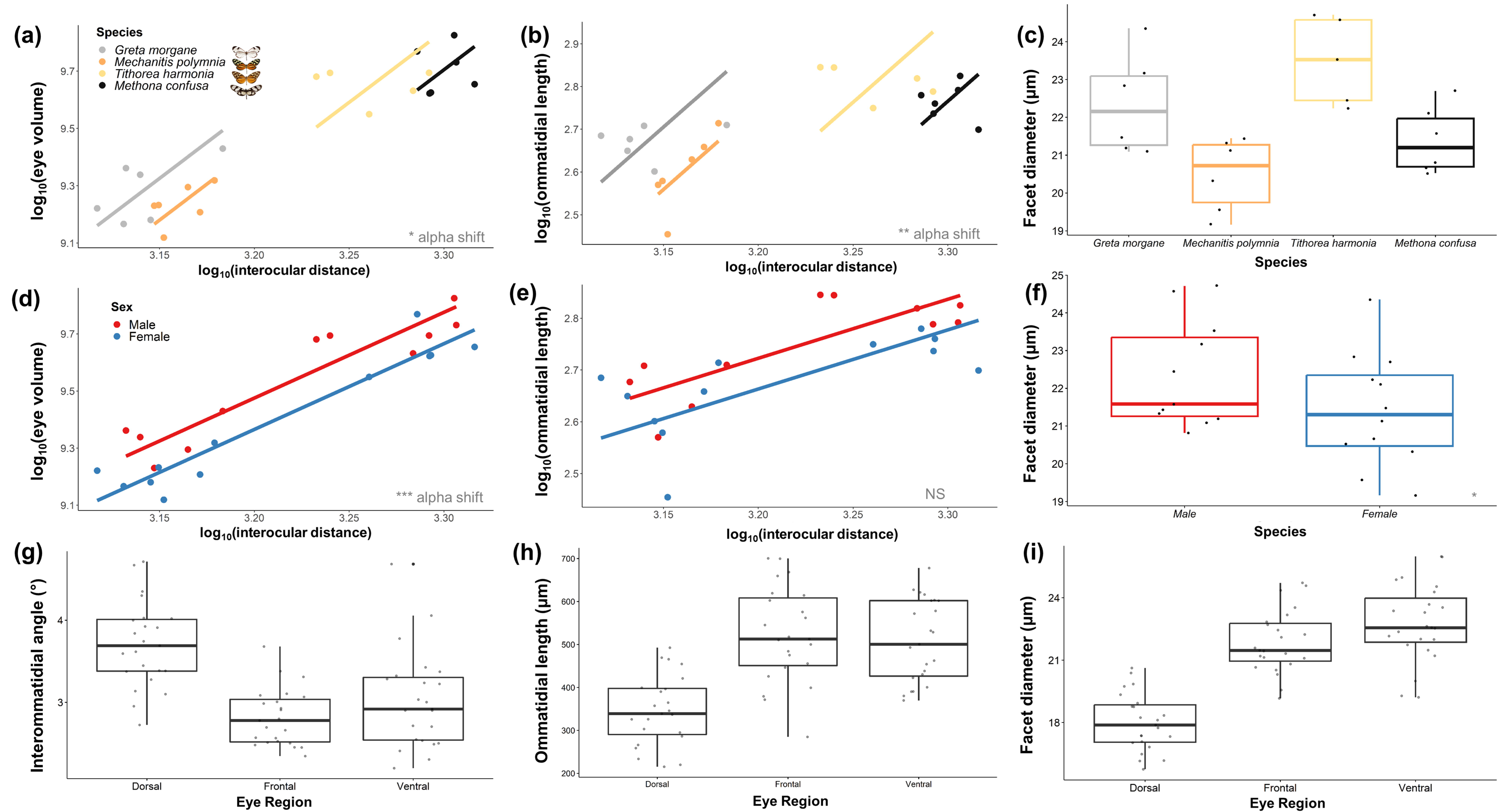
E**y**e **anatomical variation in four ithomiine species from micro-CT measurements** (n = 23, 52.17% female). (a-c) Interspecific differences in total eye volume (EYE, a), ommatidial length (*L*, b) and facet diameter (*D,* c). (d-f) Intersexual variation in EYE (d), *L* (e) and *D* (f). For EYE and *L*, log-transformed values are scaled against interocular distance (*IOD*) where an “α shift” denotes a grade shift in the relationship between these two variables from our SMATR analysis. (g-i) Intraindividual differences in interommatidial angle (ΔФ, g), *L* (h) and *D* (i) between the dorsal, frontal, and ventral eye regions. For all boxplots, medians (thick horizontal bars), interquartile ranges (boxes), values within 1.5 interquartile ranges of the box edges (whiskers) and possible outliers (data points outside the whiskers) are plotted. NS P > 0.05, *P < 0.05, **P < 0.01, ***P < 0.001.

Frontal and ventral ommatidial length (*L)* also varied between species (frontal region, F = 4.292, df = 3, p = 0.024; ventral region, F = 4.048, df = 3, p =0.029) with significant, non-allometric interspecific grade-shifts found for both eye regions (frontal, Wald χ^2^ = 23.08, df = 3, p < 0.001, (Fig. 5b; ventral, Wald χ^2^ = 15.23, df = 3, p = 0.002). For frontal ommatidia, pairwise comparisons revealed *Greta* to have relatively longer ommatidia than tiger-stripe co-mimics *Mechanitis* and *Tithorea* which showed a conserved scaling relationship (Table S6C). Interommatidial angle (ΔΦ) did not show any significant species differences when scaled against *IOD*, suggesting that variation in this trait is largely determined by variation in head size (Table S6A). Testing for differences in the absolute, untransformed values for facet diameter, *D*, revealed significant interspecific variation for the frontal and ventral eye regions (frontal, F = 8.429, df = 3, p= 0.006, (Fig. 5c; ventral, F = 6.373, df = 3, p = 0.015). Post-hoc comparisons of these models showed that the majority of significant pairwise contrasts were with *Mechanitis,* which had consistently the narrowest facets (Fig. 5c, Table S6B).

Across the four species, evidence of sexual dimorphism was observed for relative eye volume (F = 17.358, df = 1, p = 0.001) with males displaying an increased investment in eye size (Wald χ^2^ = 12.37, df = 1, p < 0.001, (Fig. 5d). However, our sample sizes were insufficient to detect whether these differences in eye volume were explained by any other measured eye anatomical traits (Fig. 5e,f). We found no interactions between sex and species for any anatomical variables.

Finally, comparing intraindividual variation in eye anatomy, we found significant variation between the dorsal, frontal, and ventral eye regions from the micro-CT scans (Fig. 5g-i).

After multiple test correction, no species, sex, or interaction effects were found, with the exception of *D* (where dorso-frontal differences between species were observed for females but not males, see Table S6E for details), suggesting these differences are mostly consistent across all individuals and species. Dorsal ommatidia were shorter and narrower than frontal and ventral ommatidia (Table S6E) and the ΔΦ of dorsal ommatidia was significantly greater than the other two eye regions (dorso-frontal, t = 8.949, df = 22, p < 0.001; dorso-ventral, t = 5.342, df = 22, p < 0.001).

### 4. Gross composition of optic lobe neuropils is generally conserved across species but there is evidence of sexual dimorphism in the presence of the ventral lobula

Staining against allatostatin revealed the retinotopic organisation of the optic lobe, particularly within the lamina and medulla where the respective cartridge and column modules are clearly visible (Fig. 6a). Distinct layers of neuronal organisation are present in the medulla, consisting of two major outer and inner divisions, separated by the serpentine layer, as seen in other ithomiines (Montgomery & Ott, 2015). Within each division are several further striations where a maximum of seven medulla layers were clearly observed in the scans of all four species (Fig. 7a). The lobula plate and the lobula have two and three layers respectively, as observed in other Lepidoptera (e.g. Hamanaka *et al*. 2012). Two smaller neuropils, the accessory medulla, found against the anteromedial edge of the medulla, and the ventral lobula, an ovoid ventromedial structure which forms a close physical association with the lobula, were more clearly visibile in whole mount anti-synapsin stains.

**Fig. 6:**
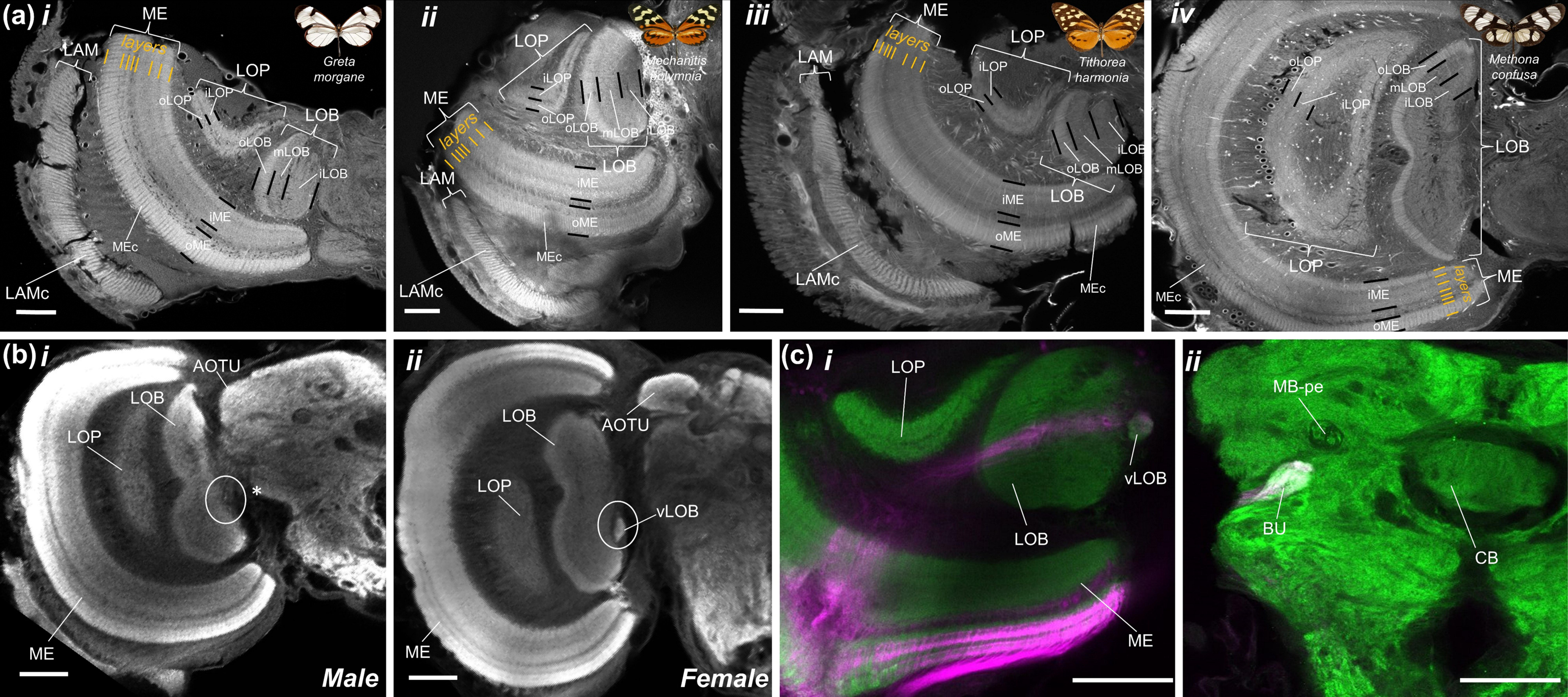
D**e**tailed **neuroanatomy from immunostained brain samples** (anti-allatostatin labelling: n = 24, 54.17% female; dextran injections: n = 24, 62.5% male). (a) Confocal images of anti-allatostatin labelled frontal vibratome sections (thickness, 80 μm) taken at approximately equivalent regions of the brain of *Greta morgane* (i), *Mechanitis polymnia* (ii), *Tithorea harmonia* (iii), and *Methona confusa* (iv). Divisions in the medulla (ME), lobula plate (LOP) and lobula (LOB) are shown using thick black lines and finer-scale layering within the medulla is shown in orange. (b-c) Sexual dimorphism in the presence/absence of the ventral lobula, a butterfly-specific optic neuropil. (b) Confocal sections of anti-synapsin labelled wholemounts of a male (i) and female (ii) brain of *Mechanitis polymnia*. (c) Confocal section of a single female *Greta morgane* brain, co-labelled against synapsin (green) and dextran injected neurons (magenta) originating from the medial optic lobe (i) and terminating in the bulb of the central brain (ii). All scale bars = 100 μm. (LAM: lamina; LAMc: lamina cartridges; ME: medulla; MEc: medulla columns; iME: inner medulla; oME: outer medulla; LOP: lobula plate; iLOP: inner lobula plate; oLOP: outer lobula plate; LOB: lobula; iLOB: inner lobula; mLOB: middle lobula; oLOB: outer lobula; vLOB: ventral lobula; AOTU: anterior optic tubercle; BU: bulb; CB: central body; MB-pe: mushroom body peduncle).

Our samples suggest that the ventral lobula, which is smaller in ithomiines than in other butterflies where it has been identified (Wainwright & Montgomery, 2022), is sexually dimorphic in *Greta* and *Mechanitis,* being totally absent in all males of these species (Fig. 6b). In *Papilio xuthus* and Heliconiini butterflies, the ventral lobula appears to act as a relay centre between the optic lobe and the mushroom body calyx (Couto et al 2023; Kinoshita *et al*. 2015). However, our dextran injections provided no evidence that the ventral lobula serves this purpose in ithomiines. The ventral lobula appears to receive inputs via the lobula, without innervation in the lobula plate (Fig. 6c), reminiscent of *Drosophila* medulla projection (Tm) neurons which receive direct synaptic inputs from photoreceptors and relay this directly to the lobula (Borst, 2009; Lin *et al*. 2015). Our dextran injections suggested the ventral lobula then projects to the bulb in the central brain, a structure which forms part of the anterior visual pathway in *Drosophila* (Hardcastle *et al*. 2021; Lovick *et al*. 2017; Omoto *et al*. 2017). Dextran staining in the anterior optic tubercle, the main optic neuropil in the central brain, was also found for some individuals, but the precise neuronal inputs to this region are not clear from our confocal scans. Injections into the mushroom body calyces also provided no evidence that these structures receive projections from the ventral lobula.

### 5. Despite a conserved composition, interspecific and intersexual variation in the visual pathway extends to investment in visual neuropils

Volumetric data from segmented micro-CT scans allowed further quantitative volumetric comparisons of the main visual neuropils (Fig. 7). Two-way ANOVAs revealed significant interspecific differences for all four structures (lamina, F = 4.617, df = 3, p = 0.019; medulla, F_3_ = 12. 543, p < 0.001; lobula plate, F_3_ = 11.806, p < 0.001; lobula, F_3_ = 14.180, p < 0.001; optic lobe, F_3_ = 11.192, p < 0.001) with no sex or interaction effects found for any neuropils. Similar to our comparisons of eye volume, allometric analysis revealed all species effects to be the result of grade-shifts in visual investment (lamina, Wald χ^2^_3_ = 17.86, p < 0.001; medulla, Wald χ^2^ = 43.42, p < 0.001; lobula plate, Wald χ^2^ = 39.27, p < 0.001; lobula, Wald χ^2^_3_ = 40.80, p < 0.001; optic lobe, Wald χ^2^_3_ = 30.68, p < 0.001; (Fig. 7b). For the medulla, lobula plate, lobula, and optic lobe as a whole, significant pairwise comparisons commonly involved *Greta*, which consistently invested more in visual neuropil, for a given brain size, than the other three species, when accounting for *IOD* (Table S6).

**Fig. 7:**
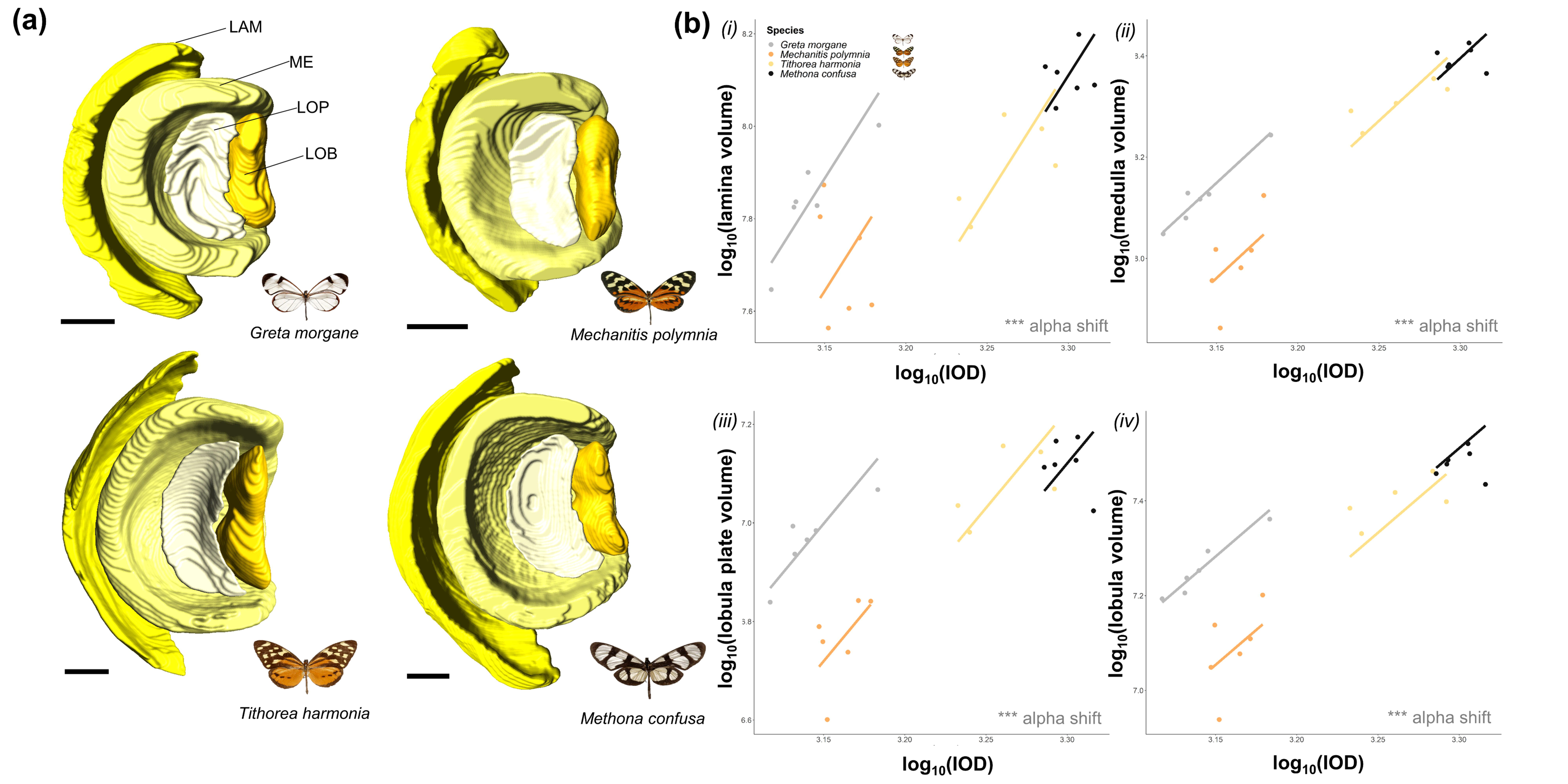
N**e**uroanatomical **variation in four ithomiine species from micro-CT measurements** (n = 23, 52.17% female). (a) Posterior surface reconstructions of the four main optic lobe neuropils (LAM: lamina, ME: medulla, LOP: lobula plate, LOB: lobula) for all four species shown to scale. Scale bar = 200 μm. (b) Interspecific scaling relationships between interocular distance (*IOD*) and the lamina (i), medulla (ii), lobula plate (iii), and lobula (iv). An “α shift” denotes a grade shift in the relationship between the two variables from our SMATR analysis. NS P > 0.05, *P < 0.05, **P < 0.01, ***P < 0.001.

A covariance matrix revealed significant correlations between the size of some of the optic neuropils and the eye (see Fig. S2). After controlling for this covariance and including species as an additional independent variable, the significant species effects remained for the eye (F = 14.232, df = 3, p < 0.001) and the lobula plate (F = 3.530, df = 3, p = 0.046).

Interspecies differences in eye size therefore appear, on the most part, to evolve in concert with investment in optic neuropils. Interestingly, interactions between functionally related neuropils appear to be obscuring sex effects as significant sex differences in the medulla were revealed after controlling for this covariance (F = 13.333, df = 1, p = 0.003) with males investing in larger medullas. This is consistent with our observed patterns of sexually dimorphic eye investment (Fig. 5d), which also remained after controlling for this covariance (F = 49.124, df = 1, p < 0.001).

## Discussion

Specialization in visual systems, likely reflects selection pressures imposed by ecological needs, which act at the functional, physiological, anatomical and molecular level (e.g. Bartholomée *et al*. 2023; Gonzalez-Bellido *et al*. 2011; Hofmann *et al*. 2009; Huber *et al*. 1997; Scales & Butler, 2016; Sugawara *et al*. 2005; Zhao *et al*. 2009). By measuring physiological, anatomical, and molecular variation in the eyes and visual neuropils of four closely butterfly species, we demonstrate multiple dimensions of variation in the visual system of Ithomiini. The 26-Myr-old adaptive radiation of these butterflies has been linked to microhabitat partitioning (Chazot *et al*. 2019; Elias *et al*. 2008; Hill, 2010), suggesting species are exposed to contrasting patterns of sensory information. By identifying aspects of the visual system that differ between species, we provide a foundation for future work linking variation in sensory ecology and sensory systems. Below we discuss our results with this eco-evolutionary framework in mind.

### Duplication and relaxed selection in UV opsins

Selection analyses on recovered visual opsin sequences revealed evidence of relaxation of selection. We found evidence of relaxed selection in the coding sequence of the UV-sensitive visual opsin in *Greta*, relative to nymphalid butterfly species living in open habitats (Fig. 2b; see Methods). Photosynthesising vegetation tends to absorb short wavelengths, particularly UV, making the light environment of the forest understory less rich in UV radiation than more exposed forest microhabitats (Endler, 1993; Théry, 2001). UV light is known to play a major role in butterfly inter/intraspecific communication and navigation (Briscoe *et al*. 2010; Brunton & Majerus, 1995; Froy *et al*. 2003; Sauman *et al*. 2005) but these cues may be less reliable for species occupying such densely shaded forest. Similar patterns were not observed for the blue and long wavelength sensitive (B and LW respectively) visual opsins. This relaxation of selective constraint might indicate a diminished role of UV-orientated behaviours in *Greta*, in comparison to its close relatives.

In contrast, we identify a duplication of the UV opsin in *Tithorea*, making this the second independent duplication of this gene identified in butterflies (McCulloch *et al*. 2017; two additional duplications of this gene have been identified in non-Papilionoid Lepidoptera but with no known functional role, see Sondhi *et al*. 2021). In *Heliconius* butterflies, which exhibit an independent UV opsin duplication, the duplicated gene experienced positive selection in some lineages but relaxed selection in others, with the latter leading to pseudogenization or down-regulated expression of this opsin in photoreceptor cells. While it is tempting to link UV duplication to increased reliance on UV cues in *Tithorea*, the duplicated copy also appears to have experienced relaxed selection, which raises questions over its functionality (Fig. 2c). Nevertheless, our data illustrate the potential for divergent repertoires of opsin genes, and selection regimes governing their evolution, across Ithomiini. Larger scale interspecific comparisons could provide opportunities to link this variation to preferences for different light environments.

### Response to temporal and spatial light variation

Despite evidence of visual opsin conservation across all four species, photoreceptor sensitivity can also be shifted by changes in eye physiology. Our eyeshine images revealed a diversity of retinal mosaic patterns, caused by fixed combinations of photosensitive rhodopsins with additional screening pigments (Briscoe, 2008; Briscoe & Bernard, 2005; Stavenga & Arikawa, 2006). These screening pigments create the red-orange reflecting ommatidia seen in our four species (Fig. 3a), providing these species with an additional spectral channel for discriminating longer wavelengths (Belušič *et al*. 2021; Zaccardi *et al*. 2006).

Intraindividual variation between dorsal, frontal, and ventral compound eye regions likely mirror spatial differences in sensory cue abundance and variability (Fig. 3A). Dorsal facets are primarily yellow-green in colour whereas frontal and ventral facets are orange-red. This indicates a specialisation for short wavelength sensitivity in dorsal ommatidia, which are exposed to a greater abundance of these wavelengths in downwelling light (Briscoe, 2008; Stalleicken *et al*. 2006). In addition, shorter ommatidia found within the dorsal region is a way of minimising self-screening for improved polarization vision, which other butterflies have been shown to utilise for orientation and navigation, particularly within the dorsal rim area (Kinoshita & Arikawa, 2014; Labhart & Meyer, 1999; Meyer & Labhart, 1992; Nilsson *et al*. 1987; Reppert *et al*. 2004) (Fig. 5h). Although the dorsal rim area was not delineated in our image data, larger interommatidial angles in the dorsal area might hint that this region is optimised for enhanced contrast detection rather than improved visual acuity. The latter would be more beneficial in the frontal and ventral regions to optimise the localisation of hostplants, foodplants and mates (Bergman *et al*. 2021; Labhart & Meyer, 1999; Labhart *et al*. 2009; Land, 1989) (Fig. 5g). A greater abundance of downwelling light hitting the dorsal ommatidia (Stavenga, 2002) could also explain the narrower facets found within this region (Fig. 5i). In our ophthalmoscopy experiment, we found that the ventral region showed longer pupillary response times which might reflect lower temporal variability in upwelling light intensity (Fig. 3d). Dorsal facing ommatidia must respond to rapid fluctuations in downwelling solar irradiance caused by gaps in the forest canopy whereas frontal ommatidia must respond quickly to changes in other types of visual contrast, such as contrast variation created by optic flow when the animal is in flight to reduce motion blur. This role in stabilising visual processing during flight potentially explains the faster response times in these eye regions (Endler, 1993; Grittner *et al*. 2022; Palermo & Theobald, 2019; Théry, 2001). However, differences in pupillary kinetics between red and yellow reflecting ommatidia, which may further interact with variation in rhabdom and photoreceptor cross-sectional area, might also explain this result.

Comparing differences in frontal pupillary response time revealed that *Greta* and *Mechanitis* show slower responses times when exposed to a bright light stimulus following dark adaptation (Fig. 3b,c). Responding more slowly to fluctuations in light conditions may reflect a history of evolution within more homogeneous environments, and a strong preference for particular light conditions. Indeed, *Greta* tends to occupy constantly shaded forest understory (Elias & Joron, 2015; Willmott & Mallet, 2004), while *Mechanitis* might consistently occupy sunlit environments and therefore respond less rapidly to changes in light intensity than those from patchier forest where *Methona* and *Tithorea* are found. In addition, the longer, thinner wings of *Methona* are indicative of greater flight speeds (Hill, 2021), which might select for faster physiological responses due to enhanced optic flow and contrast change during flight (Hill, 2021). However, we cannot rule out a degree of phylogenetic inertia in predicting these differences, given the closer relationships between these latter two species (Chazot *et al*. 2019). Larger species comparisons coupled with fine-scale ecological data (e.g. flight speed, light temporal variability, canopy cover) are required to disentangle the factors driving eye physiological variation but the well documented natural history of ithomiines (e.g. Beccaloni, 1997; Brown & Freitas, 1994; Willmott & Mallet, 2004) make them a prime model system for tackling these questions in future work.

### Interspecific differences in eye and optic lobe structure

Across our four species, we identified extensive interspecific differences in optical and neural architecture which could not be explained solely by variation in overall head size or by evolutionary relatedness. Differences were observed for relative eye size, ommatidial length, facet diameter and the relative size of all four measured optic neuropils. In contrast, the lack of interspecific differences in interommatidial angle (ΔΦ) suggest that differences in visual acuity can be fully explained by differences in overall head size (as in Jander & Jander, 2002; Rutowski *et al*. 2009).

Larger relative eye size in *Greta* could, at least partially, be explained by enhanced ommatidial length in the frontal and ventral eye regions of this species (Fig. 5a,b). Longer rhabdoms provide a greater photoreceptive surface, and longer crystalline cones would enhance focal length. Together, this would optimise the eye for increased light sensitivity at greater distances (Greiner *et al*. 2004a; Warrant & McIntyre, 1993), which may be required in low-light conditions. The large relative volume of the lamina in *Greta* is consistent with the hypothesis that this species is enhancing either the signal-to-noise ratio for improved achromatic visual sensitivity and/or the amplitude of spectral opponency for improved colour vision (Fig. 7b; Matsushita *et al*. 2022; Sterling & Laughlin, 2015; Stöckl, 2022). The lamina is retinotopically organised, with lamina monopolar cells (LMCs) within neural cartridges receiving visual input from individual ommatidia (Fig. 6a). These LMCs spatially integrate signals for enhancing light sensitivity (Greiner *et al*. 2005; Stöckl *et al*. 2016, 2020). Larger lamina volumes observed in *Greta* therefore suggests that this species devotes greater neuronal resources to processing each visual pixel. Indeed, these patterns of investment are even more extreme in the remaining three optic neuropil where *Greta* shows consistently larger visual neuropil than the other three species, controlling for head size (Fig. 7b).

Therefore, increases in relative eye size appear to positively correlate with increases in neural processing (see also Garamszegi *et al*. 2002; Corral-López *et al*. 2017).

The relatively larger eyes, optic lobes and longer ommatidia of *Greta* might have evolved to enhance visual sensitivity, reflecting the shaded low-light forest understory where this species and its co-mimics are usually found (Elias *et al*. 2008; Willmott & Mallet, 2004; Willmott *et al*. 2017). In dense forest environments, overall illuminance is likely to be lower in comparison to the more sunlit forest frequented by *Mechanitis, Tithorea,* and *Methona*, which all fly at higher elevations (Beccaloni, 1997; Elias *et al*. 2008; Endler, 1993, Hill, 2010). We note that these changes in relative investment may also arise to maintain functional visual sensitivity rather than increase it, given the constraints posed by a smaller head size (Rutowski *et al*. 2009). For example, the mean facet diameter (*D*) of *Greta* is similar to those of *Methona* and *Tithorea* despite these two species having a body size 7.12 and 4.13 times larger than that of *Greta* respectively (Fig. 5c). Hence, to overcome allometric size constraints and maintain visual performance, facet diameter must vary independently of head size.

### Sexual dimorphism in the visual system

Patterns of eye sexual dimorphism within Ithomiini match previous findings from other Lepidoptera (e.g. Everett *et al*. 2012; Meyer-Rochow & Lau, 2008; Ziemba & Rutowski, 2000) and arthropods (e.g. Zeil, 1983) where relative eye size is larger in males. Other studies have proposed this to reflect greater visual sensitivity for male-limited visual behaviours such as the searching and localisation of potential mates and territorial defence (Bergman *et al*. 2021; Everett *et al*. 2012; McClure *et al*. 2019; Pliske, 1975; Rutowski, 2000) (Fig. 5d,f). This enhanced activity may also select for males to become more visually sensitive to the presence of predators (Everett *et al*. 2012). Unlike relative eye size, sexual dimorphism was not found for structures within the optic lobe, except for the medulla but this is only revealed after controlling for covariance between optic neuropils (Fig. S2). Therefore, intersexual differences in visual ecology might promote changes in visual perception without downstream shifts in visual processing (e.g. by changing the number of lamina cartridges).

However, we also uncovered a rare case of dimorphism in the presence or absence of an apparently butterfly-specific optic neuropil – the ventral lobula (Fig. 6b). This small neuropil was absent in the ten *Greta* and seven *Mechanitis* males we examined, but present in both sexes of *Methona* and *Tithorea.* In other butterflies, the ventral lobula appears to be a relay centre directing visual projection neurons to the mushroom body calyx (Kinoshita *et al*. 2015; Couto et al. 2023). However, our own dextran neural tracing revealed no evidence of this pathway in ithomiines (Fig. 6c). Dextran fluorescence is instead detected in the bulb of the central brain which might indicate that the ithomiine ventral lobula has different functional roles which, in the case of *Greta* and *Mechanitis*, presumably coordinates female-specific behaviours. In other insects, the bulb receives projections from the anterior optic tubercle and relays these signals to the ellipsoid body of the central complex, a collection of neuropils known to coordinate navigation and orientation behaviours (Hardcastle *et al*. 2021; Heinze *et al*. 2013; Lovick *et al*. 2017; Omoto *et al*. 2017). However, the small size of the ithomiine ventral lobula in comparison to other studied butterflies suggests overall lower functional performance in this neuropil (Montgomery & Ott, 2015; Wainwright & Montgomery, 2022). Nevertheless, this work provides striking evidence of neuroanatomical sexual dimorphism, further highlighting the lability of the nervous system to vary across short ecological and temporal scales.

### Summary

Our qualitative and quantitative analysis reveals variation between and within species which potentially reflect the different ecological conditions they occupy. Previous comparative work in a wild community of ithomiine butterflies mirrors these findings, having shown that small-scale ecological shifts can predict adaptive changes in other aspects of visual neuroanatomy (Wainwright & Montgomery, 2022). Our current work further integrates information on eye structure, physiology, and molecular evolution. We present multiple lines of evidence that extensive visual system variation can exist within diverse insect radiations, hinting at the evolutionary lability of these systems to rapidly develop specialisations to distinct visual ecological niches, with selection potentially acting at both the perceptual. processing, and molecular level. We found physiological and anatomical differences between species, within species and within individuals, as well as evidence of gene duplications and relaxed selection in the visual opsins and suggest various explanatory hypotheses for these results by linking our findings with knowledge of ithomiine natural history, behaviour, and ecology. Larger comparative analyses across a greater number of species will enable the discrimination of adaptive processes from phylogenetic effects and repeated examples of microhabitat divergence and convergence within single communities make Ithomiini an ideal model system for identifying the ecological drivers of eye, brain, and visual pigment diversification in the wild.

## Acknowledgements

We are grateful to Alanna Kelly and Tom Pitman for support at the University of Bristol’s Old Park Hill Greenhouse facility, and to Amaia Alcalde for assisting in butterfly rearing. We are also thankful to the University of Bristol’s XTM Facility, Wolfson Bioimaging Facility and High Performance Computing Facility for micro-CT, confocal microscope and bioinformatics support respectively. We also thank Nick Chazot for providing the phylogeny used in Fig. 1b, Antoine Couto for his assistance with the dextran tracing and to Jessica Foley for her support with aspects of the immunohistochemistry and microscopy protocols.

## Author contributions

Conceptualisation: JBW, SHM; Methodology: JBW, EMS, EAB, FC, MJH, SHM; Validation: NWR, SHM; Formal analysis: JBW; Investigation: JBW, CS, MC, DP, EMS; Resources: EMS, EAB, FC; MJH; Data curation: JBW; Writing – original draft: JBW; Writing – review and editing: JBW, EMS, EAB, FC, MJH, NWR, SHM; Visualisation: JBW, EAB; Supervision: NWR, SHM; Project administration: JBW, SHM; Funding acquisition: JBW, SHM.

## Funding

This work was supported by the Natural Environment Research Council (GW4+ NERC DTP to JBW and a NERC IRF NE/N014936/1 to SHM), and the Air Force Research Laboratory/Air Force Office of Scientific Research (AFOSR) through the European Office of Aerospace Research and Development (EOARD) grant FA9550-18-1-7005.

### Data availability

All raw data can be found in Table S1-3 and all opsin sequence alignment and gene tree files will be uploaded to Dryad Digital Repository upon acceptance. All opsin sequences have been uploaded GenBank (accession IDs can be found in Table S1). All pseudopupil video recordings and micro-CT scans are available from Zenodo https://doi.org/10.5281/zenodo.8154379 (Wainwright *et al*. 2023).

