## Supplementary Figures for "Multiple axes of visual system diversity in Ithomiini, an ecologically diverse tribe of mimetic butterflies"

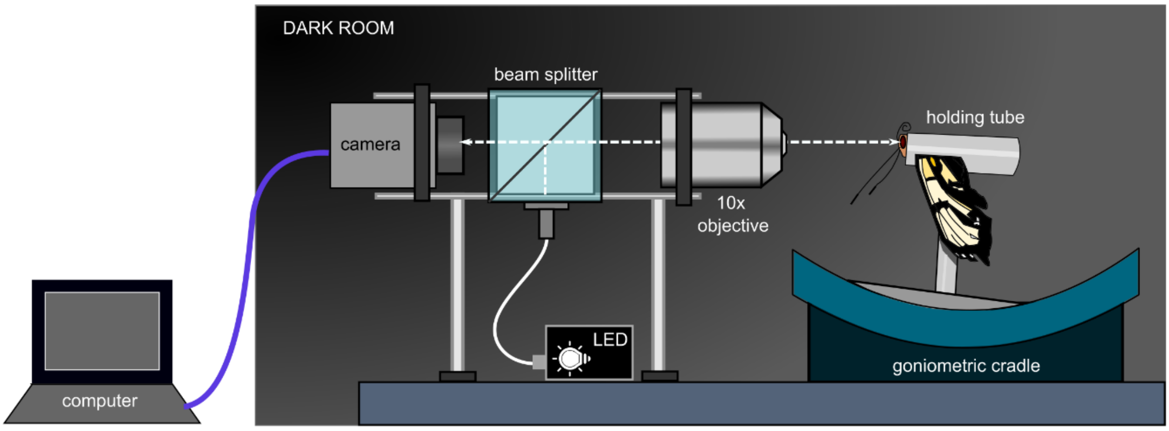


**Fig. S1: Ophthalmoscope apparatus used to image the eyeshine of live butterflies after dark adaptation.** An LED light source provided coaxial illumination to the compound eye via a beam splitter and 10x objective lens. Butterflies were mounted in holding tubes, immobilised using plasticine, and suspended at the centre of rotation of a goniometric cradle. A camera connected to a laptop with the uEye Cockpit program installed was used to record eyeshine images.

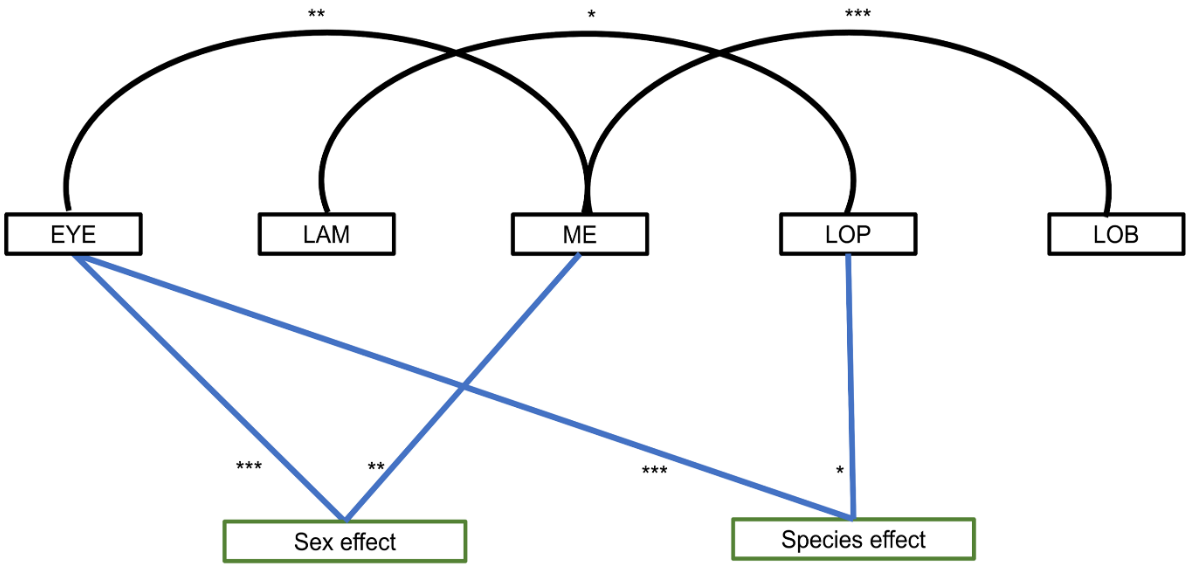


**Fig. S2:** Patterns of volumetric covariance between the four main optic lobe neuropils and the eye (EYE). Significant covariance is shown by solid black lines and structures which show significant sex/species effects after controlling for covariance are indicated by solid blue lines. NS P > 0.05, *P < 0.05, **P < 0.01, ***P < 0.001.
